# Molecular Mechanism of Capsid Disassembly in Hepatitis B Virus

**DOI:** 10.1101/2021.02.08.430262

**Authors:** Zhaleh Ghaemi, Martin Gruebele, Emad Tajkhorshid

## Abstract

The disassembly of a viral capsid leading to the release of its genetic material into the host cell is a fundamental step in viral infection. Hepatitis B virus (HBV) in particular consists of identical capsid protein monomers that dimerize and also arrange themselves into pentamers or hexamers on the capsid surface. By applying atomistic molecular dynamics simulation to an entire solvated HBV capsid subject to a uniform mechanical stress protocol, we monitor the disassembly process down to the level of individual amino acids. The strain of an external force combined with thermal fluctuations causes structurally heterogeneous cracks to appear in the HBV capsid. Unlike the expectation from purely mechanical considerations, the cracks mainly occur within and between hexameric sites, whereas pentameric sites remain largely intact. Only a small subset of the capsid protein monomers governs disassembly. These monomers are distributed across the whole capsid, but belong to regions with a high degree of collective motion that we label ‘communities’. Cross-talk within these communities is a mechanism of crack propagation leading to destabilization of the entire capsid, and eventually its disassembly. We identify specific residues whose interactions are most readily lost during disassembly: R127, I139, Y132, N136, A137, and V149 are among the hotspots at the interfaces between dimers that lie within or span hexamers, leading to dissociation. The majority of these hotspots are evolutionary conserved, indicating that they are important for disassembly by avoiding over-stabilization of capsids.

**Significance:** Hepatitis B virus (HBV) is a DNA virus that is 100 times more infectious than HIV. Despite the availability of a vaccine, the chronic infection rate of this virus is still about 300 million people globally. HBV chronic infection, for which no cure is currently available, can lead to liver cancer. Therefore, there is an unmet need to investigate the infection cycle of the virus. One of the most crucial steps in virus replication cycle is the release of its genetic material to the nucleus. During this step, the viral capsid enclosing the genetic material disassembles. However, its mechanism is unknown. Here, we utilize molecular simulations to shed light on the events leading to the capsid disassembly with atomistic detail.

## Introduction

Hepatitis B virus (HBV) chronically infects nearly 300 million people across the globe and can lead to cirrhosis, liver failure, and hepatocellular carcinoma. ^1^ Despite the availability of a vaccine, there is currently no cure for chronic HBV infection.^2^ Similar to other DNA viruses, HBV exploits the internal machinery of its host cell to replicate. To release its DNA into the nucleus, the capsid must disassemble at the appropriate time and location, and new capsids must reassemble in a very similar environment. Neither process is yet sufficiently well understood to explain this apparent hysteresis.

Unless disassembly occurs reliably, capsid contents, namely the viral genome and the viral RNA polymerase, cannot be released for replication.^1,3^ Particularly intriguing is the question of how the capsid maintains stability during its transport in the cytoplasm while on the other hand being prone to disassembly when the nucleus is reached. In-cell studies have shown that the HBV capsid disassembly occurs at the nuclear pore complex (the transport gate between the cytoplasm and the nucleus). ^4,5^ However, the details of the disassembly process remain unknown. These questions motivated us to characterize bare capsid disassembly at a molecular level. The insights we gain provide a baseline to identify the effect of genetic material on the capsid, as well as intracellular factors that control viral infection, and may guide the development of novel therapeutics^2^ or more effective synthetic nano-carrier strategies. ^6,7^

The 36-nm-diameter HBV capsid has an icosahedral symmetry composed of 240 capsid proteins (monomers) with identical sequences (Fig. 1a). Each monomer is 183 residues long and consists of two domains. The 149-residue N-terminal domain is involved in capsid assembly, while the arginine-rich C-terminal domain (residues 150-183) regulates intracellular trafficking and packaging of viral RNA.^8^ Each identical (in sequence and structure) protein monomer has a hydrophobic core that stabilizes the overall fold of its largely helical structure.^9^ The complete icosahedral capsid is composed of A sites where monomers are packed with 5-fold symmetry, and B, C, and D sites, where monomers are packed with 2-fold, 3-fold, and quasi-3-fold symmetries. ^9,10^ The five monomers around a 5-fold symmetry axis form a pentamer, and the remaining monomers form hexamers (Fig. 1). For simplicity, we refer to these arrangements simply as pentamers and hexamers without specifying their individual symmetry axes.

**Figure 1:**
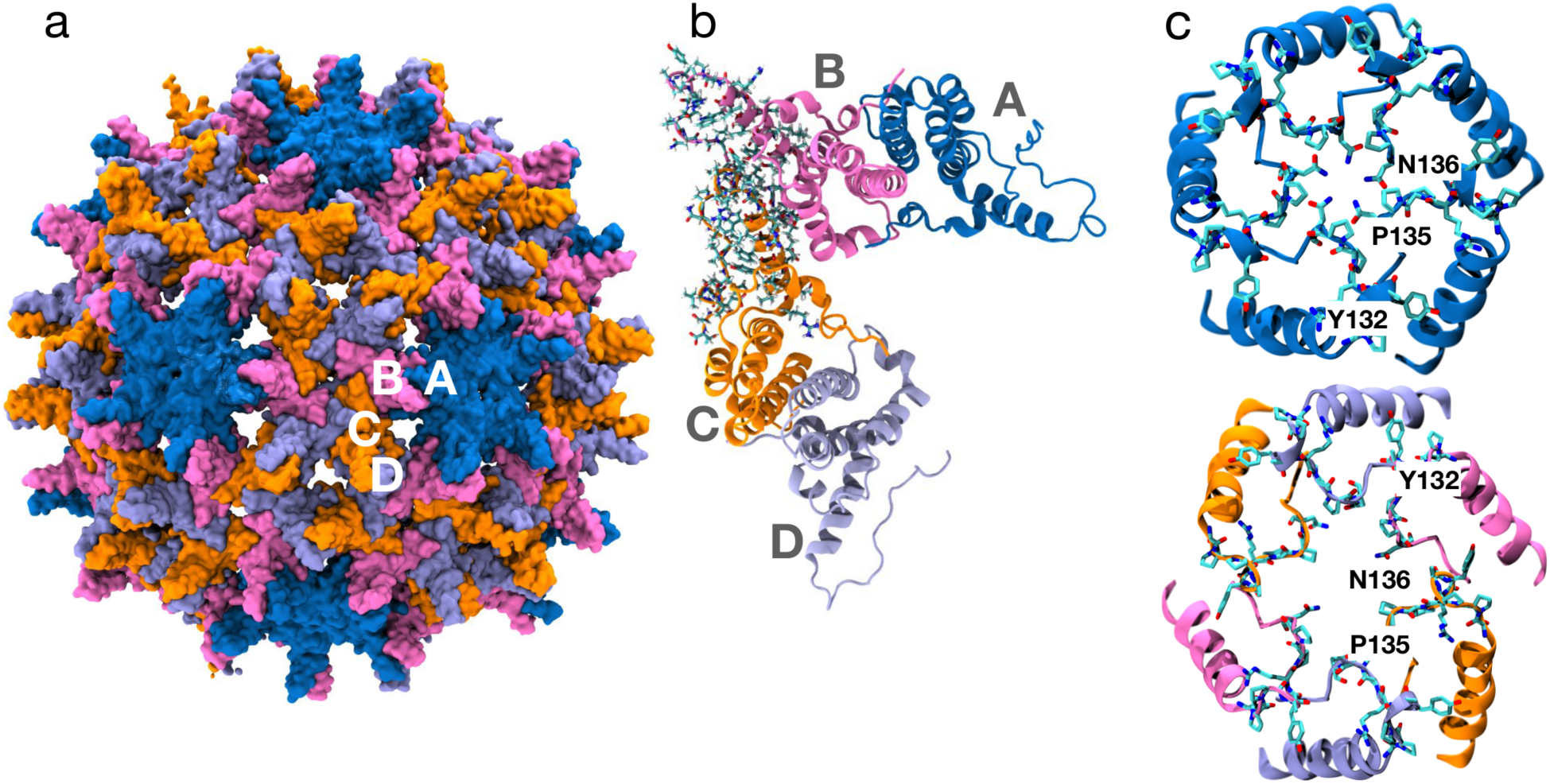
The overall architecture of the HBV capsid: a) The 36-nm HBV capsid (with a triangulation number of T = 4) is composed of four protein/monomer types (sites), A, B, C, and D, that form an icosahedral structure: site A proteins form pentamers (dark blue), while hexamers are composed of sites B, C, and D (pink, orange, and light blue, respectively). b) Examples of dimeric structures, AB and CD formed by the four sites. The capsid is formed by interactions between the dimers, with one interface highlighted here by an explicit representation of the lining side chains. c) Top view of A sites that form pentamers (in dark blue, top), and B, C, and D sites that form hexamers (bottom). The secondary structures of *α* helix 5 and the proline-rich loop (that are involved in the inter-dimer interactions) for every site are shown in cartoon representations together with some of the residues lining the openings at the 5-fold and 2-fold symmetry axes in licorice representation. While proline 135 and asparagine 136 constrict the pentamer opening, they are in different arrangements in the hexamer opening. Tyrosine 132, which is known to be crucial for the formation of inter-dimer contacts, ^9^ is shown in both oligomeric states.

Monomers form strongly bound dimers that span sites of different symmetry: AB dimers span a pentamer and a hexamer, and CD dimers span two hexamers. Within each dimer, four-helix bundles from two monomers pack against each other to form a characteristic structure that projects as a spike on the capsid surface. ^9,10^ The formation of dimers is known to be the first step in capsid assembly. Afterward, interactions formed at the inter-dimer interfaces give rise to the emergence of a stable capsid. ^11,12^

The inter-dimer contacts specifically involve an *α* helix, a proline-rich loop, and an unstructured C-terminal region of the monomers (Fig. 1b). Based solely on the degree of the surface burial of the proteins at the inter-dimer interface, the strengths of the interactions at these interfaces in pentamers and hexamers are estimated to be similar.^9,12^ However, as shown in Fig. 1c, the residues lining the openings that are formed at the symmetry axes (center) of the pentamers and hexamers and the extent of these openings are different.^9^ As a result, the biological roles of pentamers and hexamers are speculated to be different.^9^ Specifically, we anticipate that pentamers and hexamers may play distinct structural roles in capsid disassembly.

Instability and mechanical stress have been induced in capsids of several viruses, including HBV.^13–17^ Small-angle X-ray scattering experiments of cowpea chlorotic mottle virus and microsecond-long coarse-grained simulations of the triatoma virus have reported that after a rapid pH change, the capsid cracks mainly at hexameric sites, whereas pentameric structures remain intact.^14,18^ In contrast, numerical indentation simulations of phages not taking into account chemical details of interactions between capsid proteins, suggest that pentamers are more mechanically vulnerable to buckling.^19^ Similarly, atomic force microscopy (AFM) nanoindentation experiments also support buckling at pentamers. ^20^ Additionally, coarse-grained simulations coupled with AFM experiments have shown that irreversible HBV capsid deformation occurs by local shifting and bending of monomers. ^13^However, they did not report whether dimer interfaces within pentamers or hexamers are disrupted first in HBV.

Advances in computational speed and molecular dynamics (MD) force fields have made reliable atomistic MD simulations of entire viruses feasible.^21–26^ However, an atomistic simulation has not been carried out thus far to clarify how thermal fluctuations and mechanical stress combine to create structural heterogeneity during HBV capsid disassembly. Here, atomic-level MD simulations of the entire solvated HBV capsid are combined with a specifically-designed, isotropic external mechanical stress to investigate the disassembly process. We investigate the early steps of capsid disassembly, focusing on the monomer interfaces that sense and respond first to mechanical stress. We find that under stress, cracks appear on the capsid mainly between and within hexameric sites, but much less within pentamers. The chemical details of the interactions at the dimer interfaces are thus an important factor in determining the disassembly mechanism. We have also identified that within hexamers and pentamers, only a small number of monomers governs the disassembly process, pointing to the high degree of structural heterogeneity within the stressed capsid. Although these labile monomers are scattered across the capsid, they are all part of regions with a high degree of collective motions that give rise to spontaneous capsid deformation. Lastly, our simulations reveal which specific residues lining the inter-dimer interfaces are among the first ones to break their contacts. In contrast, residues within the spikes (the monomer-monomer contact within a dimer) preserve their contacts throughout the disassembly process.

## Results and Discussion

### Mechanical stress leads to the formation of cracks spanning the capsid

A viral capsid remains stable after formation, ^27^ requiring an external trigger to observe its dissociation during the simulations. We devised a spatially isotropic, external pulling potential to mechanically stress the capsid and induce disassembly (see details in Methods). Starting from a thermally equilibrated capsid (Fig. S1), the external pulling potential allowed us to observe the initiation of disassembly within affordable simulation times. We performed 20 independent disassembly simulations.

Fig. 2a shows the capsid radius as a function of simulation time reflecting the rise of the mechanical stress during the disassembly simulation. The simulations induced a maximum capsid expansion of 7%. The capsid expanded until the structure mechanically failed and cracks formed at multiple locations in the capsid (Fig. 2b). Radial expansions of ∼ 10% have been reported experimentally in viruses such as cowpea chlorotic mottle virus^28^ prior to disassembly, a result that was also reproduced by simple theoretical models.^29^

**Figure 2:**
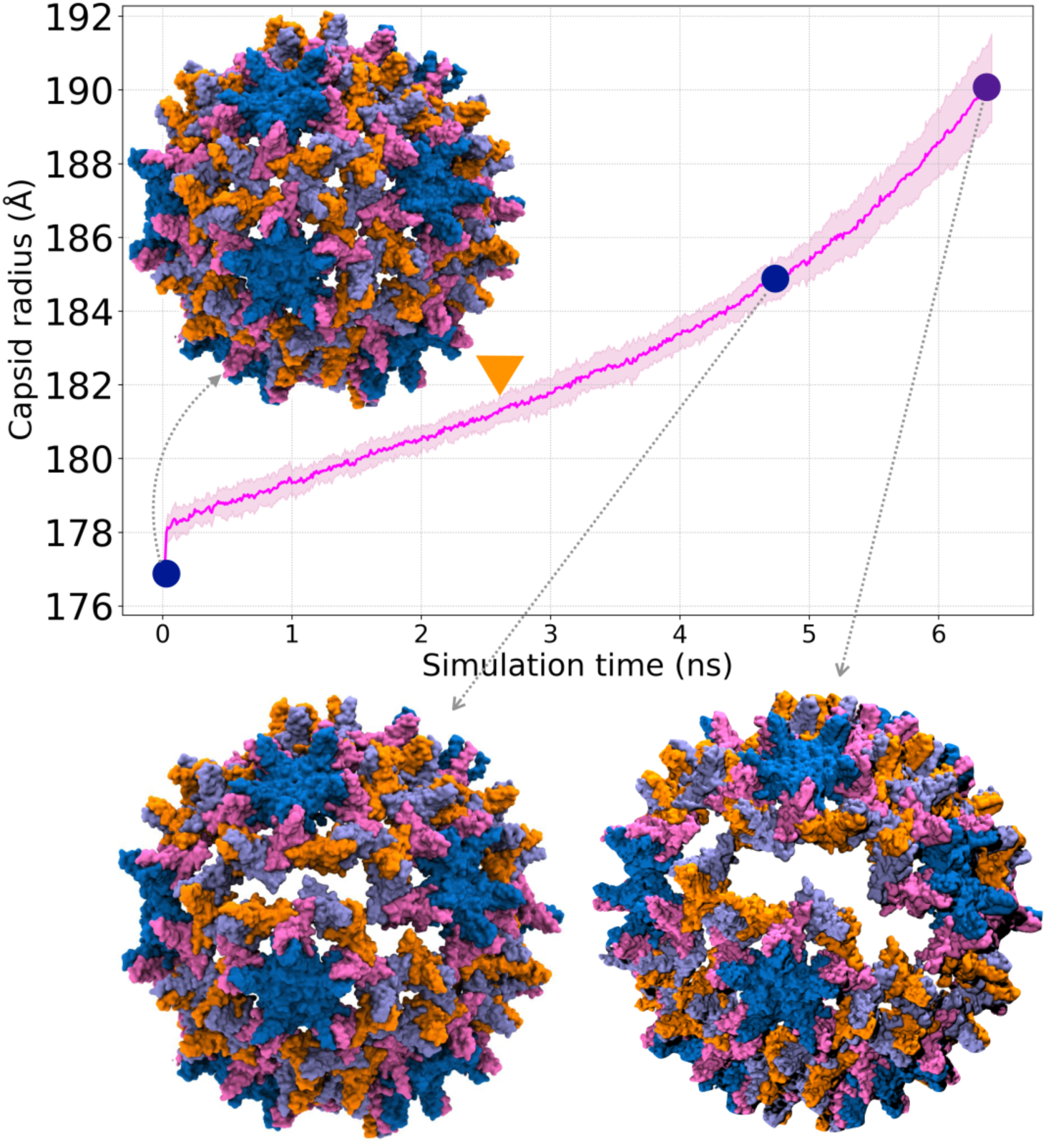
Capsid expansion under mechanical stress during the disassembly simulations: the expansion, measured as the capsid radius, was estimated by averaging the dimensions of the capsid in the three Cartesian coordinates for each simulation replica (*n* = 20). The line shows the average, and the shaded area represents the standard deviation, calculated from the 20 independent disassembly simulations. Each simulation continued until a capsid expansion of 7% was achieved. The orange triangle shows the average time for the first ‘macroscopic’ (capsid-wide) crack to form. Representative snapshots from different time points highlighting the formation of the cracks that appear in the capsid during the simulations. For clarity parts of the capsid are not shown.

We estimated the capsid expansion for the formation of the first ‘macroscopic’ crack on the capsid by averaging over all simulation replicates. A macroscopic crack was defined as a center of mass separation between neighboring monomers greater than 37 Å, which is ∼ 18% increase from the initial distance of 31.4 Å (see Analysis for details). The first macroscopic crack formed on average after the capsid reaches 2.5% of expansion. With the prescribed external potential, this corresponds to a first mean formation time of 2.6 *±* 0.3 ns from the start of the simulation (Fig. S2). By the end of the simulations (*t* ∼ 6 ns), an average of 35 macroscopic cracks per capsid are formed.

As shown in Fig. 2b, despite the isotropic external potential, there is heterogeneity in the response of different capsid proteins and the location of the cracks. These variations result from asymmetric capsid motions and distortions due to stochastic thermal fluctuations, which are also reported in previous studies of viral capsids and speculated to be of biological importance for disassembly.^22,30,31^ The variability in the average root mean square deviations (RMSDs) of the proteins within the capsid also points to their inherent structural heterogeneity due to thermal fluctuations (Fig. S3).

### Contact loss within the hexameric sites leads to crack formation

To characterize structural changes within the capsid in atomistic detail, we determined how the contacts between residues of different monomers vary during the disassembly simulations. Specifically, we developed and performed a maximum contact loss (MCL) analysis to identify monomer(s) within the capsid whose contacts with their neighbors are most reduced and therefore are considered to be the main contributors to the disassembly process (details in the Analysis section). The microscopic structural changes quantified by the MCL analysis can evolve eventually into macroscopic cracks (Fig. 2b). For every simulation frame, each monomer identified by MCL was mapped to its capsid site (i.e., A, B, C, or D). The number of occurrences of every site as MCL monomers was counted for each of the simulations individually, as shown in Fig. 3a.

**Figure 3:**
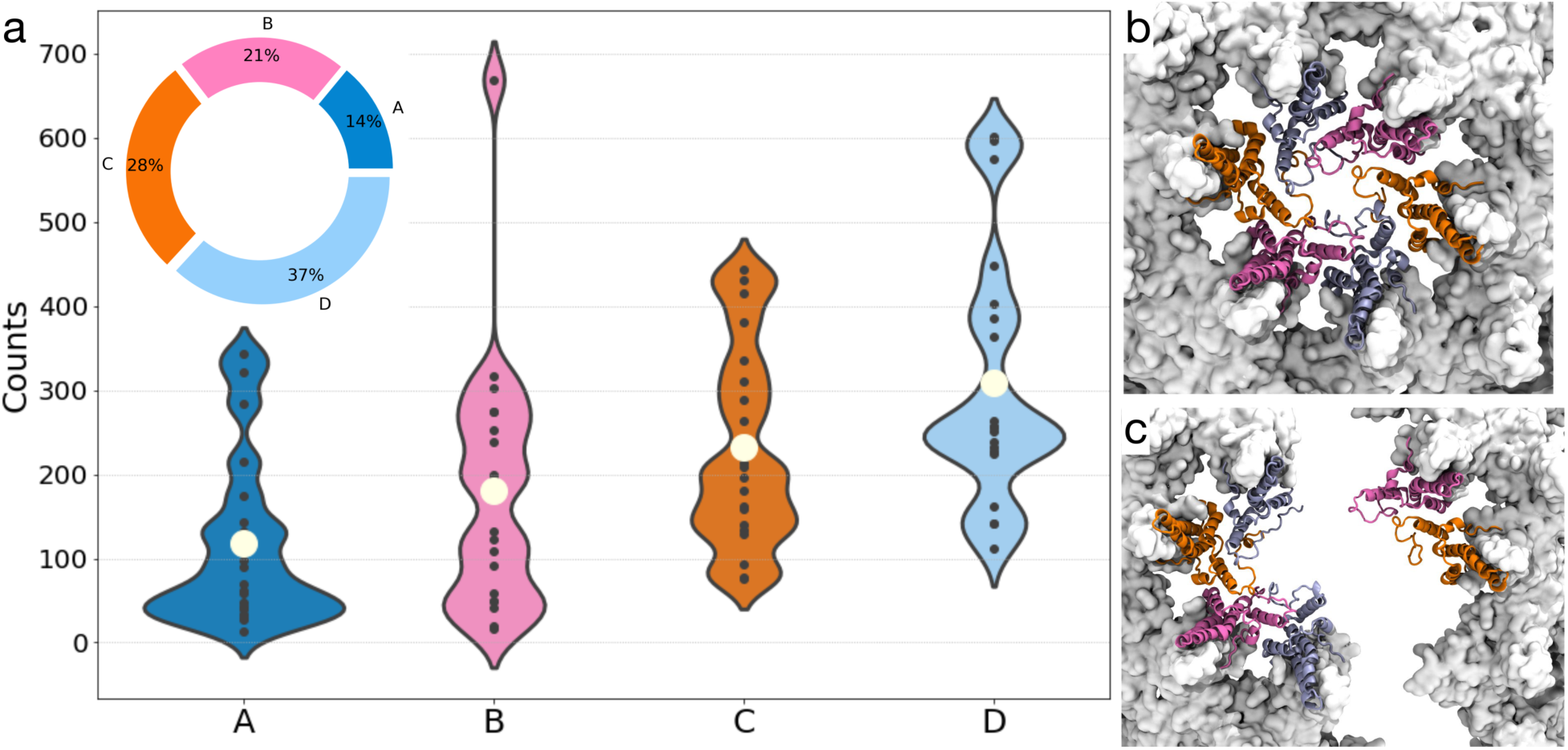
MCL (maximum contact loss) analysis of capsid proteins: a) each MCL (maximum contact loss) monomer was mapped to its site, and the occurrence of every site as MCL monomers is counted for individual simulations and shown as a violin plot. The values for all the 20 simulation replicas are shown as black dots. The average value of the counts over the 20 simulation replicas is shown as a white circle for every site. The distribution of the count values for every site represents the heterogeneity in the response of every site to the applied mechanical stress. The higher average count shows that hexamers (composed of sites B, C and D) are more involved in the disassembly process than pentamers (composed only of site A). Quantitatively, hexamers are contributing to 86% of the total monomers which are affected during the disassembly process. Snapshots of a hexameric site structure within the capsid (white surface) at the start of the simulation (b), and after the formation of a crack (c) are shown.

The MCL count provides a measure of how many monomers from a specific site were involved in capsid disassembly. Disassembly proceeds mainly through breaking of contacts at those sites that show higher counts. The broad distribution of MCL counts in all sites is in line with the structural heterogeneity during capsid disassembly that was discussed earlier. The hexamers (B, C, D sites) are the most likely units to be involved in disassembly. In contrast, pentamers participate the least in the process, as indicated by their lower average count. Of all the identified MCL monomers, 86% belong to the hexamers, and only 14% to the pentamers. This result is in contrast to the conclusions based on hydrogen-deuterium exchange mass spectrometry experiments monitoring the disassembly of virus capsids, ^15^ which could not unambiguously assign hexamers or pentamers as the most likely site of disassembly. Likewise, numerical simulations of the viral capsid based only on geometry found pentamers as the more vulnerable sites for capsid rupture. ^19^ We suggest that chemical differences between inter-dimer contacts within pentamers vs. hexamers, are responsible for favoring disassembly within hexamers first. Thus, we look at the role of individual monomers, and finally of individual residues next.

### A small number of monomers is sufficient for the progression of capsid disassembly

To identify the specific monomers (mainly within hexamers) that are key to the disassembly process, we counted the number of times each capsid monomer was classified as an MCL monomer (Fig. 4a). A higher count for a monomer signals a higher probability of its involvement in capsid disassembly.

**Figure 4:**
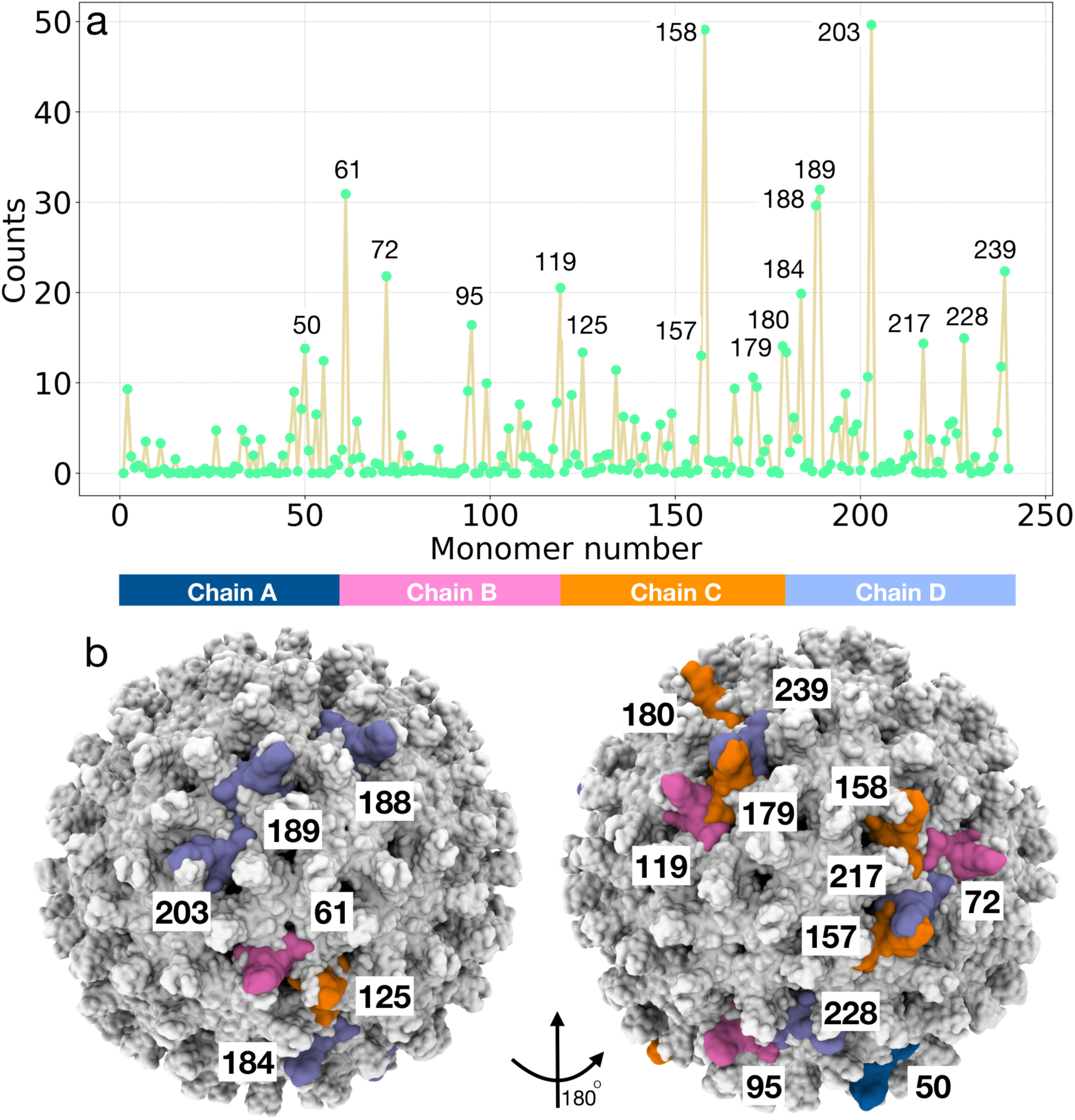
Heterogeneity in capsid proteins response to mechanical stress: a) The number of times each monomer is found to be an MCL monomer during the simulation time. Values are averaged over the 20 simulation replicas. Only 7% of monomers are classified as high-MCL (MCL count *>* 12.5) and contribute largely to the disassembly progression. These monomers are mainly from hexameric sites. b) The leading monomers, mapped onto the capsid structure, highlighting their scattered distribution across the capsid.

Monomers with an MCL count greater than 25% of the maximum value (ca. 50 in Fig. 4), account for just 17 out of the total of 240 capsid monomers. Only about 7% of monomers are directly involved in the formation of macroscopic cracks (Fig. 2b). Despite the application of an isotropic mechanical tension, the disassembly process is driven by a small fraction of the proteins due to thermal symmetry breaking between monomers. 16 of the 17 monomers appearing in Fig. 4a belong to hexameric sites, consistent with our analysis in the previous section.

To gain insight into the geometrical distribution of these monomers, we visualized them within the context of the entire capsid (Fig. 4b). The two highest MCL monomers, namely monomers 158 and 203, are positioned on opposite sides of the capsid. Each of them leads to the formation of a crack in distant parts of the capsid, suggesting that two well-separated cracks make the capsid particularly vulnerable to subsequent disassembly. The remaining 15 identified MCL monomers are distributed across the capsid, but could act cooperatively to facilitate cross-talk between distant monomers (see next section).

To examine the uniqueness of the high-MCL monomers, we varied the initial conditions of our simulations, specifically the strength of the mechanical tension and performed 10 additional simulations. These simulations, which employed a 4 times higher force, resulted in a similar overall response of the capsid, namely, the formation of cracks primarily at the hexameric sites. A different random set of monomers (Fig. 4) were identified as high-MCL monomers in these higher-force simulations (Fig. S4), but similar to the first set of simulations, only a small number of monomers (mainly within hexamers) contributed to the disassembly process.

### Collective motion of distant regions within the capsid

To study cooperativity among capsid monomers during disassembly, network analysis was performed to identify “communities” of monomers with collective motions.^32^ We define a network of capsid monomers, and calculate the Pearson correlation for their motions. A community in this network represents a collection of nodes (monomers) that move in a concerted manner with respect to each other (see Analysis for details).

The structures, the number of monomers in each community, and the composition of the site types within each community are variable among communities, demonstrating again dynamical heterogeneity within the capsid (Fig. 5a). The 17 high-MCL monomers identified above as the essential ones for disassembly are shown in the network as larger spherical nodes. These monomers mostly lie at the edges of communities, indicating that the boundaries of correlated regions are where cracking occurs.

**Figure 5:**
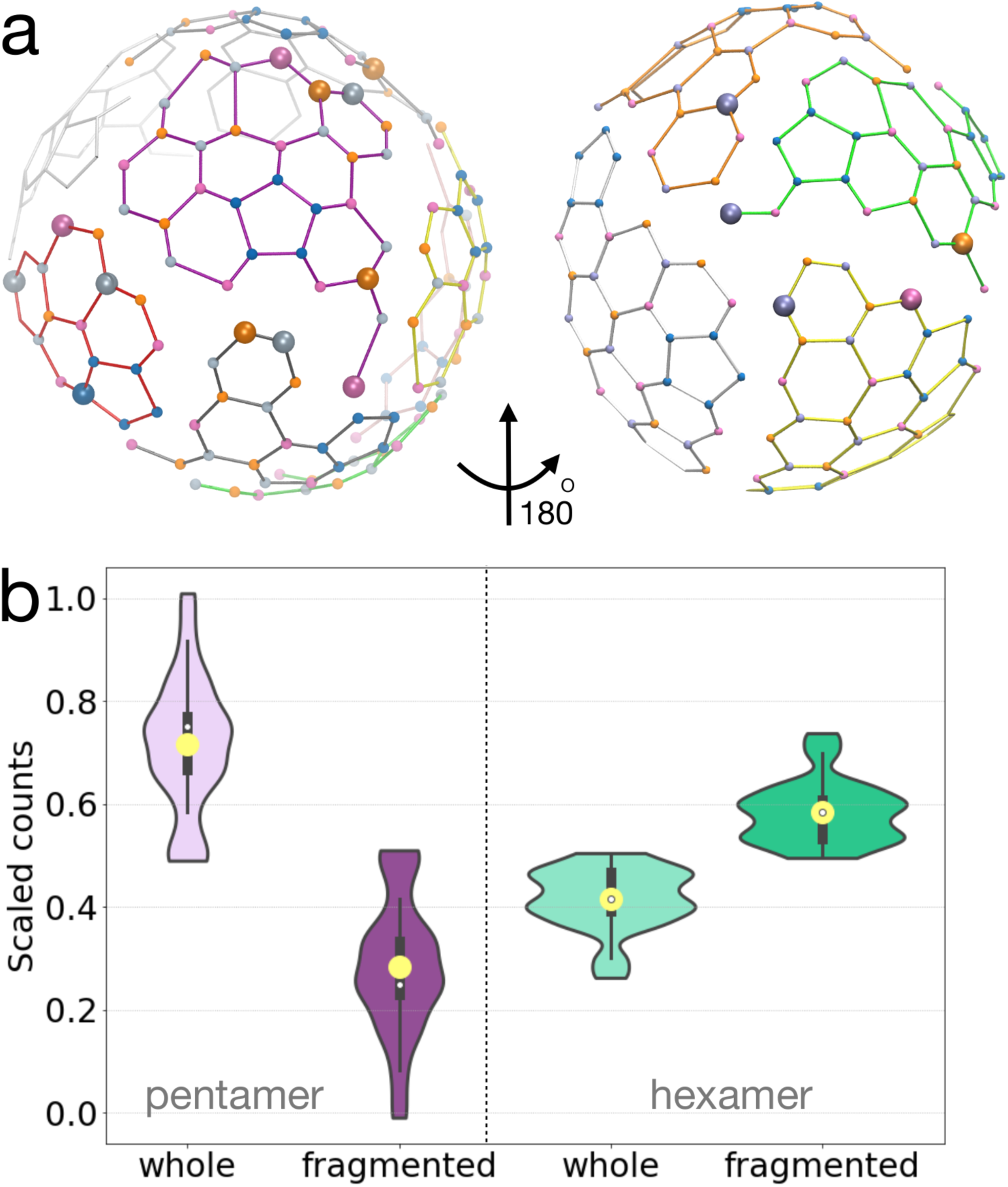
Regions within the capsid with high degrees of correlative motion: a) A network is defined using the CoM of monomers. The communities, which are the regions within the capsid network with a high degree of correlative motion, are shown on the capsid network with different colors. The high-MCL monomers are also mapped to the network (as large spheres), revealing that they either belong to one community or localized in neighboring communities. b) The pentamer and hexamer contents of communities: pentamers appear as a whole unit in 72% of communities across the capsid, whereas, hexamers only keep their unity in 42% of communities.

To establish the effects of both macroscopic cracks (Fig. 2) and contact loss between the monomers (Fig. 4) on the overall motion of pentamers and hexamers, we identified the composition of pentamers/hexamers within each community (see details in Analysis section). As shown in Fig. 5b, the pentamers keep their integrity and only 28% do not lie entirely within a single community. In contrast, 58% of hexamers are split over more than one community. During disassembly, the dynamics of monomers in about 9 out of 12 pentamers are more correlated with one another than to other monomers, whereas the same is true for only about 12 out of 30 hexamers.

### Specific residues located at the inter-dimer interfaces are hotspots of capsid disassembly

We now turn our attention to residue-level analysis to determine the specific residues that are involved in capsid disassembly (details in Analysis). In Fig. 6a, a high count means that a residue loses its contacts with neighboring monomers with a high probability. Hence, the residues showing the highest counts are the hotspots for crack formation within a monomer. Examining the secondary structure of a monomer (bottom of Fig. 6a), one can see that these hotspots are located in helices *α*1, *α*2, and *α*5, as well as in the proline-rich (P) loop and the C-terminal region. Helix *α*5, P-rich loop, and the C-terminal region are known to be involved in the inter-dimer interactions,^33^ where the sum of many weak interactions contributes to the stability of the entire capsid structure. ^27^

**Figure 6:**
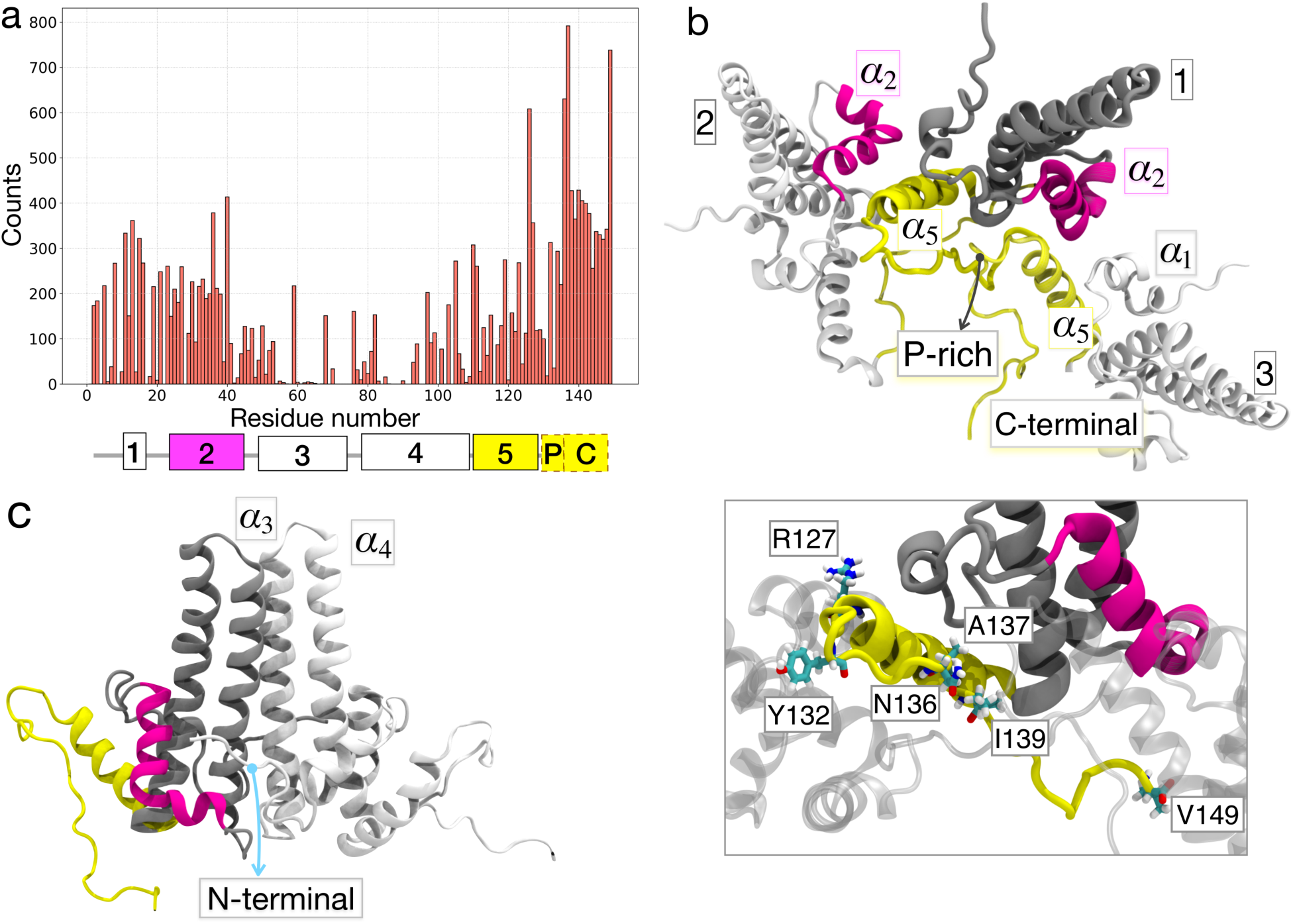
Disassembly hotspots: a) Counts calculated as the number of times a specific residue loses its contacts with neighboring monomers during the disassembly. Disassembly hotspots are located at the *α*1, *α*2, *α*5, P-rich loop, and the C-terminal region of the proteins, as indicated by the secondary structure of the protein segments. b) Each dimer form two inter-dimer interactions: dimer-1 (colored in gray, yellow, and magenta) shown in top view, together with two neighboring dimers (numbered 2 and 3) that form inter-dimer interactions with it. For clarity, instead of the entire dimers, only the interacting monomers are presented. The *α*5 and *α*2 helices of dimer-1 interact with *α*2 and *α*5 of dimers-2 and -3, respectively. The *α*5 helix, P-rich and C-terminal are colored in yellow, and *α*2 is shown in magenta. The close-up view shows the selected hotspots that were highlighted in the text. c) Side view of a dimer, shown with two helices of *α*3 and *α*4, that are involved in the intra-dimer interactions.

To set the stage for discussing the hotspot residues, we need to describe the inter-dimer interfaces in more detail. As shown in Fig. 6b, each dimer has two inter-dimer interfaces: its *α*5 helix, as well as the *α*2 helix, interact with two different dimers. Therefore, *α*2 is also involved in the inter-dimer interfaces. Additionally, the presence of hydrophobic and electrostatic interactions between *α*1 of one dimer with *α*2 of another (Fig. S5), suggests that helix *α*1 can also contribute to the inter-dimer interactions. The P-rich loop and the C-terminal region are involved in both interfaces.

Among the disassembly hotspots, N136, A137, and V149 are involved in both of the inter-dimer interfaces (Fig. 6b close-up view). In addition, residues R127, I139, and Y132 are known to be important for capsid assembly. ^9^ Most of these hotspot residues (Fig. 6a) are conserved among known HBV sequences. ^9^

In contrast to disassembly hotspots, are the residues that retain their contacts (“tight-contact” residues) to their neighboring monomers throughout disassembly. In Fig. 6a such residues have low count values because their contacts with the neighboring monomer are not affected during disassembly. The tight-contact residues are mainly localized in the *α*3 and *α*4 helices, which both lie at the intra-dimer interface (Fig. 6c) and are involved in the formation of dimers (AB or CD): helix *α*3 is associated with extensive interactions between the two dimer-forming monomers and is responsible for holding them together. Despite the fact that *α*4 does not directly contribute to dimer formation, it is a hallmark of dimer stability and the HBV capsid spikes. ^9^ The presence of tight contacts within these helices suggests that during disassembly dimers (AB or CD) keep their quaternary structure and remain bound, whereas inter-dimer interfaces could be classified as quinary structure. ^34,35^ These findings are in accordance with the small-angle X-ray scattering experiments performed after changing the pH of the capsid environment. It was observed that dimers, but not monomers, are the final product of a fully dissociated capsid. ^14^

## Conclusion

Our findings shed light on the *in vivo* capsid disassembly mechanism. It was debated whether capsid disassembly occurs in the cytoplasm or at the nuclear pore complex (NPC).^36^ The current experimental evidence suggests that the latter scenario is more likely. ^5,37^ Specifically, the intrinsically disordered regions in the pore of the NPC called phenylalanine-glycine (FG) repeats were identified as a possible site for interacting with the capsid, that can induce instabilities.^5^ From a recent structure of the NPC,^38^ it can be deduced that the capsid may be surrounded by the FG repeats from all directions. Therefore, the spatial arrangement of the external potential we applied on the capsid (see details in Methods), can resemble the tugging effect of FG repeats in the NPC. Hence, we propose the hypothesis that capsid disassembly at the NPCs can also be initiated at mainly the hexameric sites, by breaking contacts at the inter-dimer interfaces.

In conclusion, using an external pulling potential together with MD simulations we studied the initiation of the disassembly process of an entire HBV capsid down to the single residue level. We find that structurally heterogeneous cracks appear on the capsid, mainly within the hexameric sites (sites B, C and D). We have identified a small number of capsid protein monomers that govern this crucial step of the virus life cycle, by developing a maximum contact loss (MCL) analysis. The MCL monomers are found to be distributed far from each other across the capsid but connected by regions with high degree of collective motions within the capsid. Lastly, we have identified residues that are hotspots for disassembly. They are located at the interfaces between dimers, whereas the contacts between monomers within a single dimer remain stable.

These results advance our understanding of a key step in viral infection, which may guide the design of novel pharmacological interventions against this major health concern.

## Materials and Methods

### System preparation and molecular dynamics simulations

We constructed an HBV capsid model using the most complete crystal structure with 3.95 Å resolution^10^ (PDB ID: 2G33) as shown in Fig. 1. The atomic structures of site A included 148 residues, sites B and C, had 147 residues, and site D, contained 146 residues. To simulate the 149-N terminal domain, the unresolved residues at each site were added using psfgen. The Solvate plugin of VMD^39^ was used to solvate the capsid with TIP3P water molecules. The water padding around the capsid was chosen large enough to ensure a distance of at least 40 Å between periodic copies. After neutralizing the system by Na^+^ ions using the VMD Ionize, 0.15 M NaCl was added to the water box. The CHARMM36m force field^40^ was used for the proteins. The determination of the protonation states was done with PropKa.^41^ The solvated simulation box included approximately 6.5 million atoms and box dimensions of 407 *×* 404 *×* 402 Å^3^.

The system was then minimized in four steps: 1) 15,000 steps with all heavy atoms restrained; 2) 15,000 steps with only protein heavy atoms and ions restrained; 3) 30,000 steps with protein heavy atoms restrained, all with a restraining potential constant of 5 k *cal/mol/* Å^2^; and 4) 30,000 steps with no positional restraints. Then, we increased the system temperature from to 298.15 K in 6 ns at a rate of 25 K per 500 ps. Finally, the system was equilibrated as an NPT ensemble using the Langevin thermostat (damping coefficient of 5 ps^−1^) and Nosé-Hoover Langevin piston method (200 fs piston period and 100 fs piston decay).^42,43^ Periodic boundary conditions and a timestep of 2 fs were used. Long-range electrostatic forces were computed by the particle-mesh Ewald (PME) algorithm. ^44,45^ Lennard-Jones interactions were cut off at a distance of 12 Å. All simulations were performed with NAMD 2.13 and 2.14. ^46,47^ After about 150 ns of MD simulations, the root mean square deviation (RMSD) plateaus (Fig. S1), and the system reaches equilibrium. We performed an additional 250 ns of MD simulation (total time = 400 ns) and capsid coordinates taken at 350, 360, 370, 380, 390, and 400 ns of the equilibration simulation were used as initial structures for the disassembly simulations, as described below.

### Application of mechanical stress with an external potential

We intend to simulate the initiation of disassembly without fully breaking the capsid into dimers. To induce capsid disassembly under mechanical stress, we used an isotropic pulling potential using grid Forces,^48^ as implemented in NAMD.^47^ Such isotropic potentials can actually be applied to proteins even experimentally. ^49^ Briefly, an arbitrary potential can be defined on discrete grid points, interacting with specified atoms within the simulation system. ^48^ After interpolating a continuous potential function from the defined potential, the forces on the atoms are determined. As shown in Fig. 7, we designed the grid potential to start from the interior of the capsid expanding to the boundaries of the simulation box. This design allows us to create a radial pulling force according to the potential:

**Figure 7:**
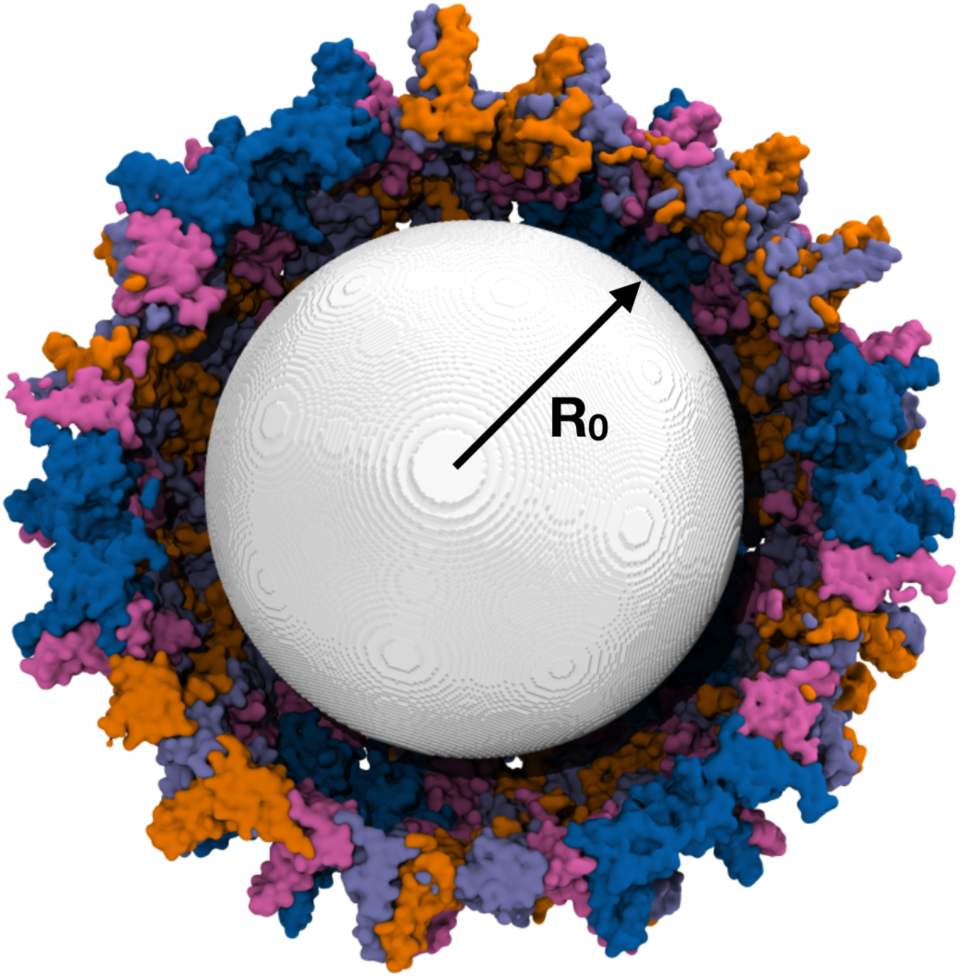
The spatial arrangement of the external potential: The onset of the isotropic external potential (R_0_) is shown in a white surface in a cross section of the capsid. The applied potential extends to the boundaries of the simulation box.

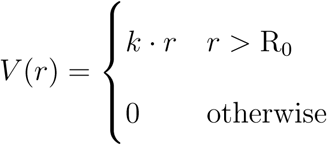

where, R_0_ is the onset of potential at 100 Å, a distance chosen to guarantee that all capsid atoms are affected by the potential, and k = −2.5 *×* 10^−4^ k*cal/mol/*Å is the force constant. We increased the force constant k to higher values in additional simulations, but qualitatively similar results were obtained (see Fig. S4). Grid points were separated by 1 Å.

To improve sampling, we performed 20 independent disassembly simulations with our devised pulling potential. Each simulation was stopped as soon as the radius of the capsid reached 189.5 Å. This value was chosen to ensure that initiation of capsid disassembly was observed, but at least a distance of 14 Å between periodic images of the capsid was maintained.

## Analysis

### Formation of macroscopic cracks in the capsid

To characterize the order of the ‘macroscopic’ (capsid-wide) cracks formed, we identify the protein monomers involved in the cracks and the time of their appearance. To this end, we calculated the distance between the centers of masses (CoMs) of the neighboring monomers during the simulation. A crack is considered to form when the CoM distance between the two neighboring monomers becomes larger than 37 Å, from an initial distance of 31.4 Å. Examination of the trajectories shows that this distance threshold ensures that the transiently rebound events are not counted, and therefore, neighboring monomers have permanently lost contact.

In an intact capsid, there are 360 CoM contacts between neighboring protein monomers among the 240 monomers. As a crack starts to form in the capsid, the CoM distance between two protein monomers will exceed the threshold and hence the total number of contacts within a defined threshold decreases by one from the initial value of 360. Therefore, monitoring the reduction in the number of contacts and the monomers associated with these contact losses, will inform us about the formation and progression of cracks within the capsid during the simulation. The results of this analysis is presented in Fig. S2.

### Maximum contact-loss (MCL) analysis

To identify the monomers that are most affected by capsid disassembly, we performed a maximum contact loss (MCL) analysis on the obtained MD trajectories. Specifically, we first identified the residues contacting any monomer, by considering all residues within a distance of 4.5 Å from it. Then we determined how many of the residues contacting each monomer at the beginning of the simulation lose their contacts at a later time. Finally, for every simulation frame, we identify the monomers that have lost the most contacts with their neighboring monomers. We call these the maximum contact loss or “MCL” monomers.

### Network and community analysis

A network is constructed on a set of nodes which are connected by edges. These nodes were defined as the CoMs of the individual capsid monomers. The edges are placed between pairs of nodes if any heavy atoms from the two corresponding monomers are within 35 Å of each other for at least 75% of the trajectory. Each edge is weighted by the weight *W*_*ij*_ = −*log*(|*C*_*ij*_|), where *C*_*ij*_ are the Pearson correlation value for the motion of the two nodes. On the constructed network, we identified the communities within the complex. The community detection follows Girvan–Newman algorithm, and they represent the nodes that are densely interconnected with each other.^32^

### Pentamers and hexamers composition of communities

By determining the fraction of pentamers and hexamers within each community, we aim at identifying the dynamical aspects of these oligomeric units. To this end, first we simply counted the number of pentamers or hexamers that are enclosed within each community or are fragmented between different communities. Then to compare the values for pentameric and hexameric units, the obtained values were normalized by the total number of pentamers and hexamers in the capsid (12 and 30, respectively). The obtained data together with the average values (averaged over the data from 20 simulation replicas) are presented in Fig. 5b.

### Identifying disassembly hotspots

The MCL monomers report on the proteins leading the disassembly. We use these monomers to identify the residues that have vulnerable contacts (hotspots) during disassembly. Specifically, for every MCL monomer, we determine the number of times each residue from a neighboring monomer loses contacts with it as a result of crack formation. The same calculation was repeated for every simulation replica and averaged over all replicas.

## Acknowledgements

The research presented in this study was supported by grants from NIH (P41-GM104601 to ET and R01 GM093318 to MG). The simulations were supported by the resources provided by the Blue Waters at UIUC and Comet-XSEDE through grants ILL bban and TG-MCB180022 (to MG and ZG), respectively.

## Notes

### Competing Interest Statement

The authors have declared no competing interest.

