## Supplemental figure for "Molecular Mechanism of Capsid Disassembly in Hepatitis B Virus"

### Supplementary Information
### Molecular Mechanism of Capsid Disassembly in Hepatitis B Virus

Zhaleh Ghaemi[a,b], Martin Gruebele[a,b,c,d,*], and Emad Tajkhorshid[a,b,c,e,*]

[a]*Department of Chemistry,* [b]*Beckman Institute for Advanced Science and Technology,*
[c]*Center for Biophysics and Quantitative Biology,* [d]*Department of Physics, and*
[e]*Department of Biochemistry, University of Illinois at Urbana-Champaign*

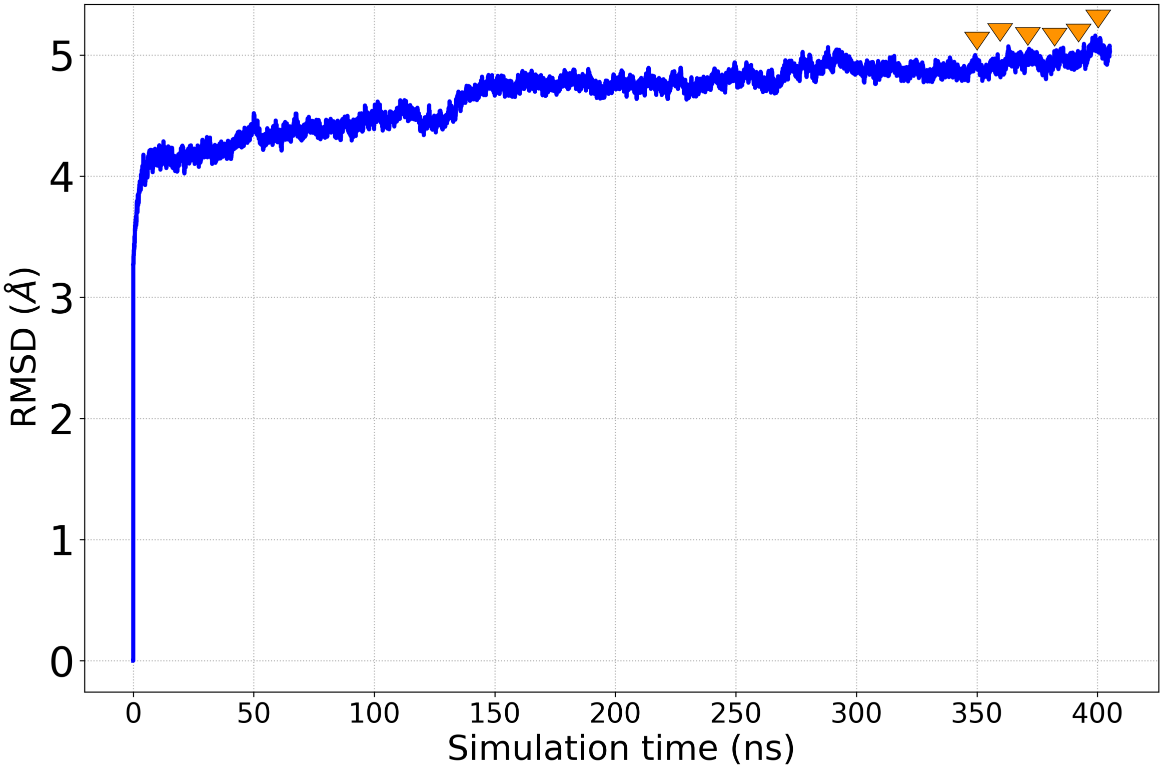


Figure S1: **RMSD of equilibrium simulation of the capsid**: The root mean square deviation (RMSD) of capsid during the equilibrium simulation. Frames at 350, 360, 370, 380, 390, and 400 ns from this equilibrium simulation (marked with orange triangles) were selected as the starting points for the disassembly simulations with the external potential, in order to average out any possible effect of the initial frame on capsid response to the external potential.

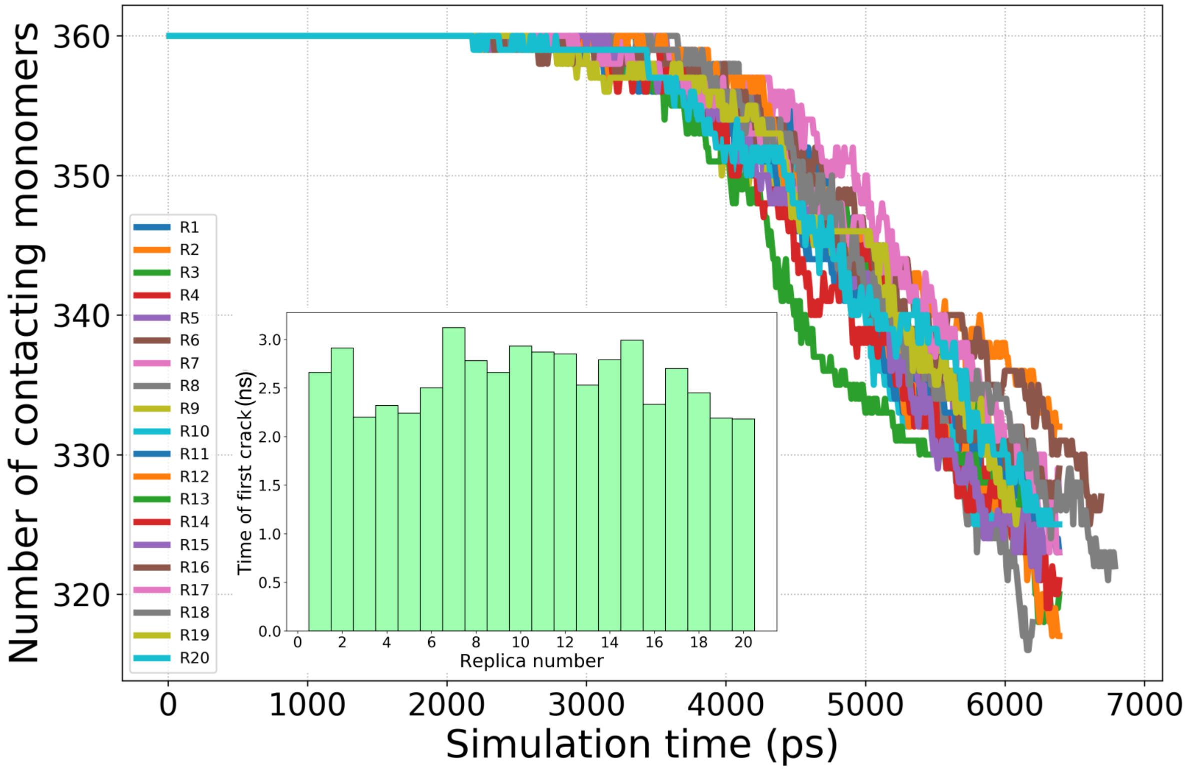


Figure S2: **Formation of macroscopic cracks in the capsid during the disassembly simulations**: As the simulation progresses cracks form in the capsid, namely the CoM distances become greater than the threshold of 37 Å. Therefore, the number of contacting monomer pairs decreases from the initial value of 360. The total number of contacting protein neighbors versus time for different simulation replicas are shown. At the end of the simulations, on the average, 35 cracks have been formed in the capsid, and it takes $2.6 \pm 0.3$ ns from the start of the simulation to observe the first crack, as shown in the inset. Inset shows the time points at which the first crack forms in all 20 simulation replicas.

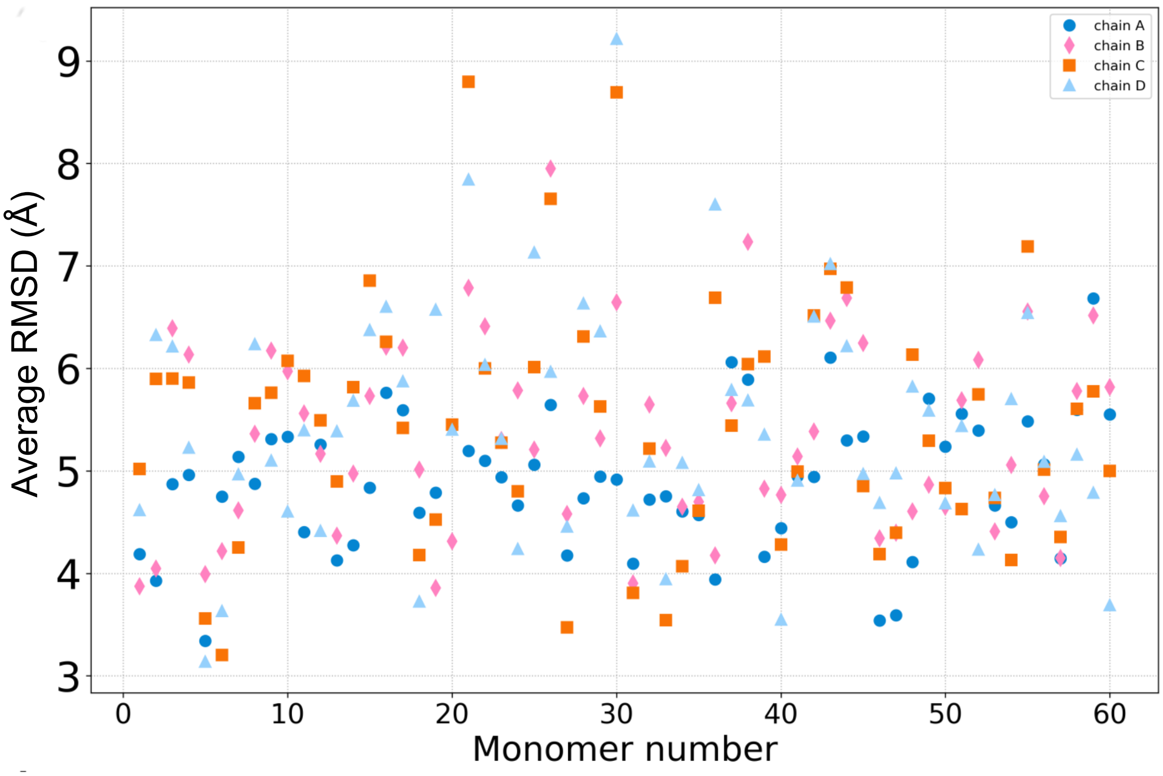


Figure S3: **RMSD of the capsid during simulations with applied external potential**: a) The RMSD of each site averaged over the simulation time when alignment was performed over the backbone atoms of the entire capsid. There are 60 monomers of each site, therefore, the horizontal axis range is from 1 to 60. The variations within the average RMSD indicates the heterogeneities within the capsid.

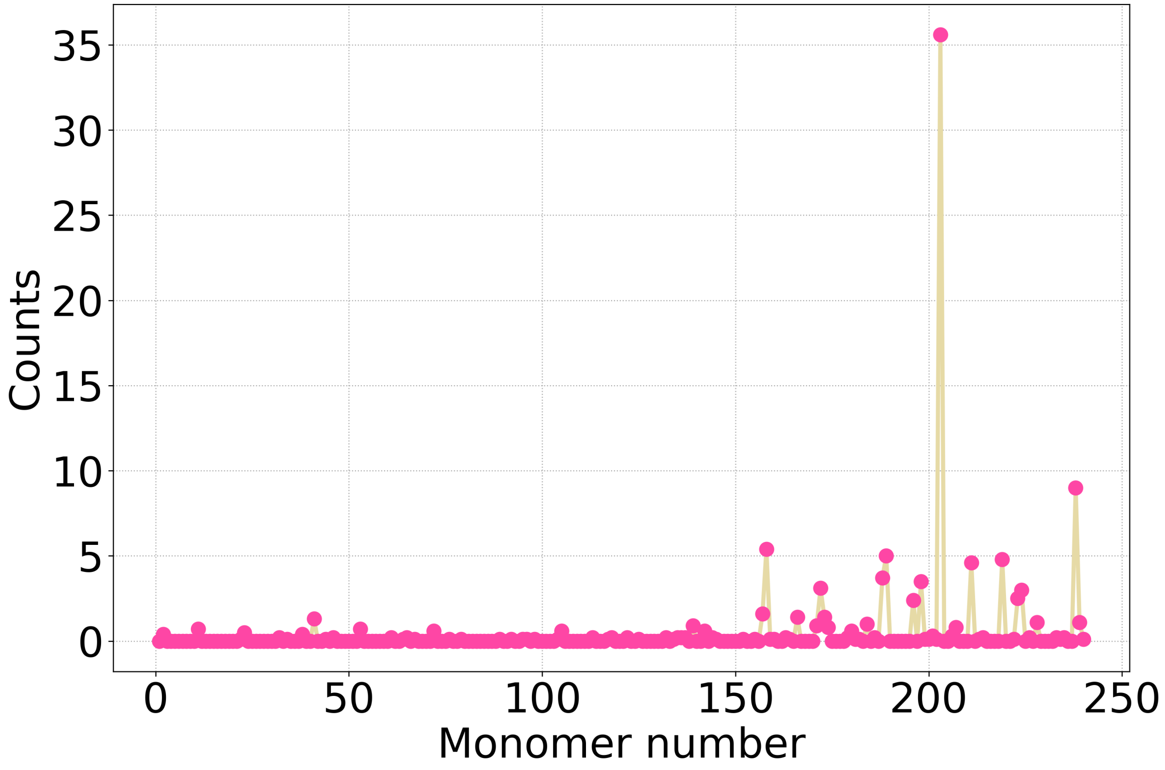


Figure S4: **Monomers that are leading the progression of disassembly within simulations with higher force.** The number of times each monomer is found to be an MCL monomer during the simulation time. Values are averaged over the 10 simulation replicas. These simulations were performed under 4 times higher force than the ones presented in the main text.

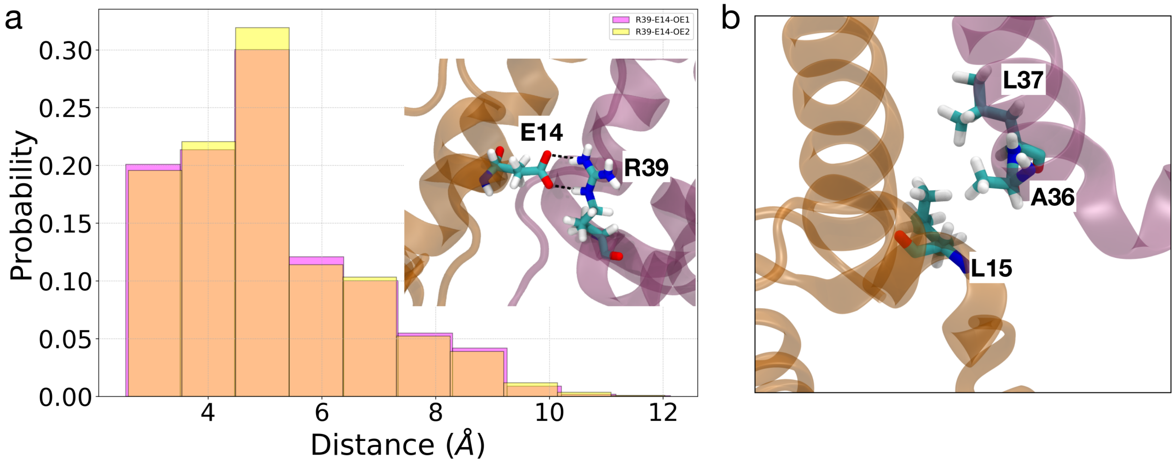


Figure S5: **The electrostatic and hydrophobic interactions between the $\alpha$1 and $\alpha$2 helices at the inter-dimer interfaces:** a) Histogram of two distances between the side chains of R39 (of $\alpha$2) and E14 (of $\alpha$1) during the 250 ns of equilibrium simulation. The range of the distances shows the presence of hydrogen bond and electrostatic interactions between these two residues. b) Residues L15 (of $\alpha$1), A36 and L37 (of $\alpha$2), contribute to hydrophobic interactions at the inter-dimer interfaces.
